# Integrating temporal genomic and transcriptomic analyses to decipher genetic basis of feed intake in dairy cattle

**DOI:** 10.64898/2026.02.07.704532

**Authors:** Caelinn James, Lingzhao Fang, Zhiguang Wu, Jayne Hope, Mike Coffey, Bingjie Li

## Abstract

**Background:** Food intake is a complex trait in living organisms, where the genetics of food intake have been widely studied in humans, mice, Drosophila, cattle, pigs, chicken, and fish. In dairy cattle, intake of feed is highly linked to individuals’ energy balance, health, production, efficiency, and the environmental footprint of the individual to the society. Recent studies have provided solid evidence of the genetic variation of feed intake (FI) in dairy cattle population, but the genetic basis and molecular mechanism of dairy feed intake is still far from clear especially considering the lactation cycles of dairy cattle. This study aims to integrate stage-dependent genome-wide association (GWA) analyses, regional heritability mapping (RHM), and RNA-seq gene expression analyses to identify temporal functional variants associated with cattle dry matter intake (DMI) across multiple stages in lactation cycles. A total of 750,000 daily DMI records from 7,500 lactations of 2,300 cows were available with animals’ genotype and pedigree information. Total RNA-seq from blood were generated for 121 individuals in this population from 2 lactation stages. Data were split into multiple lactations stages for GWA, RHM, and transcriptomic analyses.

**Results:** Stage-dependent GWAS and RHM identified 21 significant loci associated with DMI across multiple lactation stages. A total of 45 candidate genes were identified from GWA and RHM. Among all the 45 genes, six genes were later found significantly differently expressed between high and low feed intake animal groups using gene expression information from RNA-seq data. These genes show links to sugar and adipose metabolism, milk production, body weight, dopamine-reward pathways and immune functions.

**Conclusions:** Our multi-omics analyses provide molecular evidence that the genetic basis of cattle DMI across lactation is not static. Temporal genomic variants associated with FI were identified with their transcriptomic patterns investigated, decoding the molecular mechanisms underlying DMI. Overall, the associated variants and candidate genes uncovered herein decoded genetic architecture of dairy feed intake on a temporal and multi-omics basis, enhancing the understanding of basic biology of dairy feed intake and informing breeding strategies aimed at improving dairy feed efficiency.

## Introduction

Food intake is a fundamental fitness trait in living organisms as it is essential for maintenance of energy homeostasis and survival [1, 2]. The genetics of food intake have been widely studied in humans [3, 4], mice [5, 6], Drosophila [7], cattle [8, 9], pigs [10, 11], chicken [12, 13], and fish [14, 15]. In livestock species like cattle, intake of feed is highly linked to individuals’ energy balance, health, production, efficiency, and the environmental footprint of the individual to the society [16–19]. Recent studies have provided solid evidence of the genetic variation of feed intake in dairy cattle populations worldwide [20–22] and initiated genomic selection to genetically improve feed efficiency in dairy populations [23–26]. However, the genetic basis of dairy feed intake is still far from being understood. Understanding the genetic basis of feed intake improves understanding of the cattle energy mechanism and provides functional genomic information that benefit cattle precision breeding by providing proper weights into prediction for complex traits like feed efficiency.

The genetic complexity of feed intake in dairy cattle is largely due to lactation cycles [27]. It has been widely observed that dairy cattle perform differently between different stages of lactation in production and feed intake [18]. For first lactation only, it has been reported that the heritability for feed intake varies between 0.1- 0.5 across lactation stages [21], with low genetic correlation between feed intake in early and mid-late lactation [21]. A few genome-wide association (GWA) analyses have been conducted to identify quantitative trait loci (QTL) associated with dairy feed intake, focusing on individuals during first lactation [28] or individuals in mid-lactation stage across parities [29–31], but very few significant signals were identified partly due to limitation in power. The limited proportion of genetic variance explained by single markers for feed intake from previous GWA analyses reflects the limited power of single SNP analyses to detect causal variants of small effects for dairy feed intake.

For complex traits like dairy feed intake, the identification and validation of small-effect QTL by genomic analyses have proven challenging by GWA alone. Regional Heritability Mapping (RHM) provides an estimate of the regional heritability attributable to a small genomic region [32], and has the power to detect regions containing multiple alleles that individually contribute too little variance to be detectable by GWAS as well as regions with single common GWAS-detectable SNPs [33–36]. In addition to genomic analyses, integrating omics data such as transcriptomic information offers the possibility to validate functional variants identified from genomic analyses for complex traits like feed intake, providing effective validation and deeper insights of the molecular mechanisms underlying complex traits like feed intake.

Previous GWA studies on dairy feed intake have mostly focused on first lactation animals or animals in mid-lactation stage when multiple-lactation data were used. Pooling data roughly across lactations and stages can lead to decreased power of genomic analyses for feed intake considering the differences in genetic basis for feed intake across parities [18] and across lactation stages [37, 38]. Meanwhile, the genetic basis of feed intake in early lactation, which differs significantly from other stages and closely links to cattle health and fertility, was rarely studied. Taking all these potential changes in genetic basis into account, time-dependent genetic analyses (including time-dependent GWA and RHM) are useful and essential to gain a better understanding of the genetic mechanisms underlying feed intake over time.

In this study, we aimed to integrate time-dependent GWAS, RHM-based fine-mapping approaches, and gene expression transcriptomic analyses to (1) investigate the change of genetic basis of dairy feed intake over time using genomic and transcriptomic information; (2) identify genetic variants associated with feed intake in multiple lactations and at essential lactation stages including early lactation through genomic and transcriptomic analyses.

## Materials and methods

### Phenotypic data

The study builds on 50-year dairy recording on feed intake, milk production and composition, body weight and condition, health status, and reproductive events per animal at SRUC’s Dairy Research Centre, Scotland, UK. The research centre was based at Langhill herd, Edinburgh from 1970s to September 2001, and was subsequently transferred to Crichton Royal Farm in Dumfries, Scotland. Over time, a longitudinal database has been built-up for 7,400 lactations of 2,471 Holstein cows (Lactation 1 to 4) for a total of over 2 million daily records on feed intake, production, conformation, health, and fertility (until October 2024).

Feed was offered in individual feed bins (HOKO-system, Insentec B.V), and feed offered to cows and feed refusals were measured individually to calculate the feed intake per cow. The dry matter content (DM%) of the feed was recorded and aligned with feed intake records to obtain daily dry matter intake (DMI) (kg/d) per individual. A total of 701,457 daily DMI records were obtained for 2,101 Holstein cows from lactations 1-4 (until October 2024). The DMI records below 0kg or higher than 3.5 standard deviations were discarded in the phenotype quality control. The daily DMI records were then averaged on a weekly basis to obtain average daily DMI (kg/d) per animal per week, focusing on the first 44 lactation weeks (corresponding to 305 days in milk) for each lactation. After filtering, a total of 175,245 daily DMI records averaged on a weekly basis (kg/d DMI per week per lactation) from 2,099 Holstein cows remained in the dataset for analyses. The lactation curve for DMI across lactations and weeks are illustrated in Figure 1.

**Figure 1.**
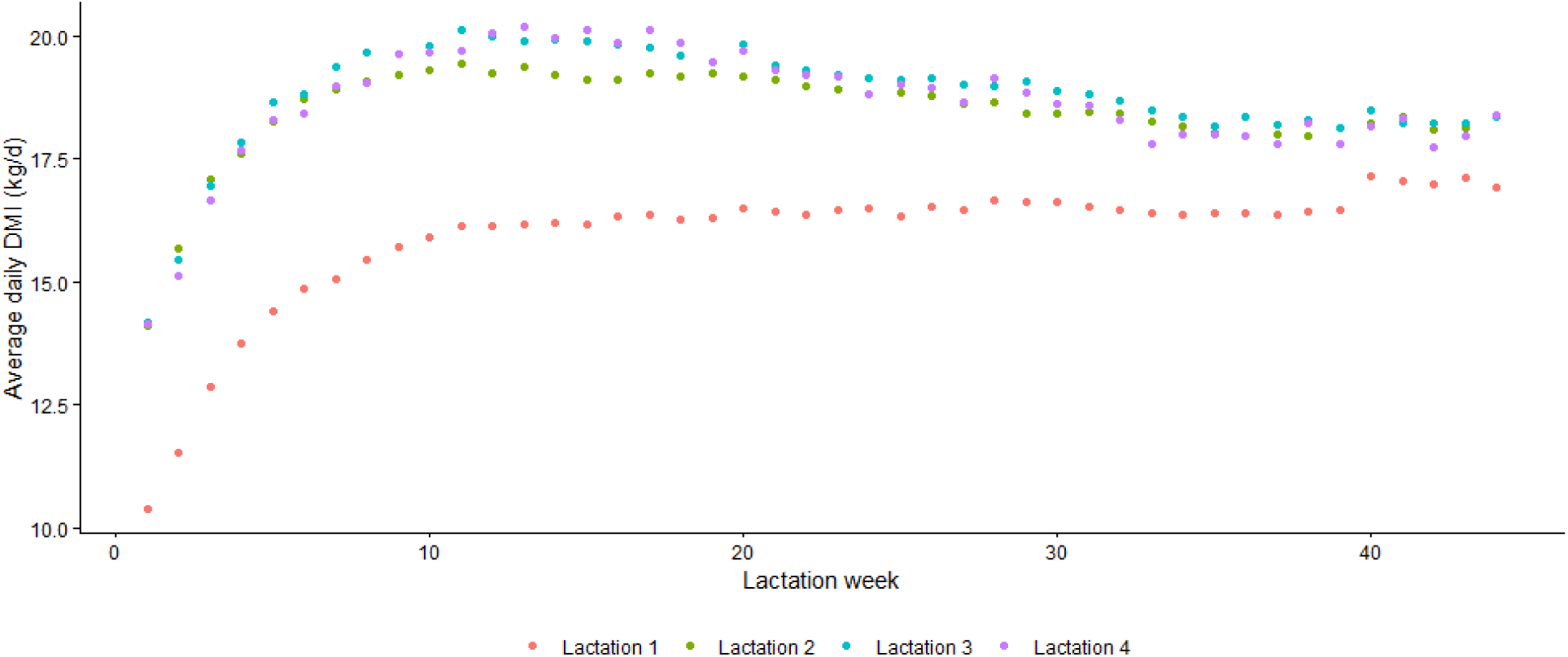
Average daily dry matter intake (DMI, kg/d) over the first 44 lactation weeks across Lactation 1 to Lactation 4.

### Genotypic data

The 80K imputed genotypes for the studied animals were extracted from the UK national database for genomic evaluation in dairy cattle. The 80K SNP panel was used in the UK national genomic evaluation for dairy cattle and developed based on original Illumina Bovine 50K BeadChip, 777K HD BeadChip and several other commercial genotyping chips plus extra gene tests and large-effect sequence variants [39]. In the quality control process, animals and SNP with call rates <0.90, SNP with minor allele frequency <0.05, monomorphic SNP, SNP deviating from Hardy-Weinberg equilibrium expectation, and animals with parent-progeny Mendelian conflicts were removed from the data set. After data filtering, 76,930 SNPs were retained in the genotypes. The locations of the SNPs were mapped to the bovine reference genome ARS-UCD1.3.

### Trait Definition

Considering the differences between lactation stages in the phenotypic level and genetic basis of DMI, the overall dataset was divided into four subsets based on parities and lactation stages: (1) Primiparous cows, early lactation (Early Lactation 1); (2) Primiparous cows, mid-late lactation (Mid-Late Lactation 1); (3) Multiparous cows, early lactation (Early Lactation 2-4); (4) Multiparous cows, mid-late lactation (Mid-Late Lactation 2-4). Early lactation stages contain data from lactation week 1-8 (corresponding to the first 56 days in milk), and mid-late lactation stage contains data from lactation week 9-44 (corresponding to the 57-305 days in milk). The four periods were taken as four traits in the following analyses where all following analyses were carried out for each of the four periods separately. The number of genotyped and phenotyped animals are shown in Table 1.

**Table 1.** The number of genotyped and phenotyped individuals, and the number of records of dry matter intake (DMI), for primiparous cows (Lactation 1) and multiparous cows (Lactation 2-4) at early lactation stage and mid-late lactation stage.

| Lactation | Lactation Stage | Number of records | Number of individuals |
| --- | --- | --- | --- |
| Lactation 1 | Early lactation<sup>1</sup> | 10,171 | 1,447 |
|  | Mid-late lactation<sup>2</sup> | 43,714 | 1,449 |
|  | Total | 53,885 | 1,484 |
| Lactations 2-4 | Early lactation<sup>1</sup> | 20,198 | 1,201 |
|  | Mid-late lactation<sup>2</sup> | 78,981 | 1,159 |
|  | Total | 99,179 | 1,513 |

<sup>1</sup>Early lactation stage includes lactation weeks 1-8
<sup>2</sup>Mid-late lactation stage includes lactation weeks 9-44.

### Phenotype pre-correction

To account for the differing number of weeks in each stage, phenotypes were pre-corrected for the environmental effects of year/season, age, feed group, parity, permanent environment and week of lactation using PREDICTF90 from the BLUPF90+ software suite [40, 41] and the following model:

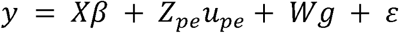

where *y* is the vector of phenotypic values for DMI; *X* is a design matrix linking individual records with the vector of fixed effects β (year/season, age, parity, week of lactation and feed group), *Z_r_* is an incidence matrix that relates the random effect of permanent environment to the individual records; *u_pe_* is the associated vector of non-genetic permanent environment effect; *g* is the vector of additive genetic random effects with *W* the incidence matrix; and ε is the vector of residuals. It is assumed that var(g) ∼ *N*(0, *G*σ_g_^2^), where σ *^2^*is the additive genetic variance and *G* is the genomic relationship matrix.

The phenotypes were pre-corrected (adjusted) to include only the additive genetic effect and the residual for each record, and the mean phenotype was calculated for each individual.

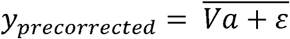

This pre-correction was performed separately for each lactation stage.

### Genome-wide association studies (GWAS)

We performed GWAS for DMI in 4 stages: (1) DMI in early lactation stage in primiparous cows (Lactation 1); (2) DMI in mid-late lactation stage in in primiparous cows (Lactation 1); (3) DMI in early lactation stage in multiparous cows (Lactation 2-4); (4) DMI in mid-late lactation stage in multiparous cows (Lactation 2-4). The DMI at these four stages were treated as independent traits considering their potential difference in genetic basis. Pre-corrected DMI for these 4 stages from earlier phenotype correction were used as the phenotype for GWAS using GCTA [42] and the *–mlma-loco* option:

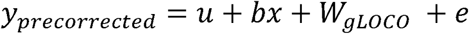

where *y_pre-corrected_* is the phenotype for DMI for each stage; *u* is the mean term; *b* is the marker effect (fixed effect) of each SNP to be tested for association, *x* is the SNP genotype indicator variable coded as 0, 1 or 2; *gLOCO* is the vector of additive genetic random effects excluding the chromosome on which the focal SNP resides with *W* the incidence matrix. It is assumed that var(g) ∼ *N*(0, Gσ_g_^2^), where σ *^2^*is the additive genetic variance and *G* is the genomic relationship matrix generated by GCTA using the Yang method [43]; *e* is the residual. To account for inflated p-values caused by use of the LOCO genomic relationship matrix (GRM), we adjusted the p-values by the lambda factor [44].

In the significance test for markers’ effects, Bonferroni correction was used to correct for multiple testing with the genome-wide significance threshold being p-value = 0.05/76,930=6.5e^-07^ (i.e., 0.05/the number of tested markers) and the genome-wide suggestive threshold being p-value = 1/76,930=1.23e^-05^ (i.e., 1/the number of tested markers).

To test whether multiple suggestive SNPs in the same loci were independent of one another, we performed conditional analysis; The genotypes of the SNP with the smallest association p value from each associated region were added to the GWAS model as a fixed covariate and removed from the GRMs and genotype data. The GWAS analysis was then re-run accounting for those associations.

### Regional heritability mapping (RHM)

We performed regional heritability mapping (RHM) [32] using GCTA [42] using the pre-corrected phenotypes for DMI in each of the 4 lactation stages. The genome was split into 50-SNP sliding windows (mean length of 1.6Mb per window) with a 25-SNP overlap. We calculated the log likelihood ratio test (LRT) statistic for each genomic window in a RHM model compared against the null model (the model with no regional genomic component fitted). The models were as follows:

RHM:

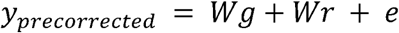

Null model:

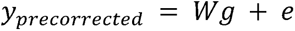

where *y_precorrected_* is the pre-corrected phenotype for DMI, *g* is the vector of the random whole-genome additive genetic effect with *W* as the incidence matrix, following var(g) ∼ *N*(0, *G*σ_g_^2^) where σ*_g_^2^* is the additive genetic variance and *G* is the genomic relationship matrix estimated using markers from the whole genome; *r* is the vector of the random regional additive genetic effect with *W* the incidence matrix, following r ∼ *N*(0, G_r_σ_r_^2^) where σ*_r_^2^* is the regional additive genetic variance explained by the markers in each window and *G_r_* is the genomic relationship matrix estimated using markers in the window; *e* is the vector of residuals. The GRMs were generated by GCTA using the Yang method [43].

In the significance test, we considered a window to be significantly contributing to the trait variance if the resulting LRT p-value is smaller than the significance threshold of 0.05/2986=1.7e^-05^ (i.e., 0.05/the number of tested windows), or suggestive if the p value is smaller than the suggestive significance threshold of 1/2986=3.3e^-04^ (i.e., 1/number of tested windows).

Any significant windows were then subdivided into 10-SNP sliding windows with a 5-SNP overlap, and RHM was performed again. In the case of consecutive 50-SNP windows being suggestive, they were first merged into one larger window before being split into 10-SNP windows. We then determined which 10-SNP windows were significantly contributing to trait variance by defining the significance threshold as 0.05 divided by the number of 10-SNP windows for each lactation stage respectively.

Because linkage disequilibrium is extensive across the cattle genome, neighbouring windows can be correlated and a window in LD with one overlapping a causal variant may itself exceed the significance threshold. To resolve whether significant signals on a chromosome reflected multiple independent causal regions, we applied an iterative conditional RHM procedure. For each chromosome with significant windows, we first identified the 10l1SNP window that explained the highest proportion of genetic variance and rel1ran RHM for the remaining significant 10l1SNP windows while fitting that window’s GRM as an additional random effect alongside the wholel1genome GRM and the regional GRM for the tested window. Importantly, if multiple 10l1SNP windows remained significant after a conditional step, we selected from those remaining the window that still explained the highest proportion of genetic variance and fitted that window’s GRM as the next random effect. This process was repeated iteratively, always adding the GRM of the most influential window that remained significant after conditioning, until no 10l1SNP windows on the chromosome remained significant.

### Generation of gene expression profiles for the study population

Total RNAs were exacted from the whole blood of 121 phenotyped cows on farm in the study population in August 2023 using Tempus™ Blood RNA Tube with Tempus Spin RNA Isolation Kit (Thermo Fisher Scientific, Cat. No. 4342792 and Cat. No.4380204) according to the manufacturer’s instructions to obtain high-quality blood RNA profiles for individual cow samples. The extracted blood RNA were sequenced for total RNA-sequencing using BGI sequencer in October 2023. The Fastp software [45] was used to check the read quality of the RNA-seq and to filter out any low-quality reads. The reads were then mapped to the reference genome (ARS-UCD1.3) using STAR [46], and the results were sorted and converted to BAM format using SAMtools [47]. Duplicate sequences were marked by Picard [48], and gene expression quantification was performed using featureCounts [49]. Raw counts were transformed into transcripts per million (TPM), and genes with a TPM of less than 0.1 were excluded from further analysis. This resulted in a total of 24,753 genes with expression in at least one individual, of which 18,542 are protein coding.

The gene expression quantification for each gene was pre-corrected for feed group, lactation number, age and lactation stage using the model.matrix function in R [50].

In March 2024, 90 of the 121 individuals had total RNA extracted again. The RNA was processed using the same methods and techniques as the batch from August 2023.

### Validation of DMI candidate genes by gene expression

The closest gene located in the vicinity of significant SNPs within 0.5Mb upstream and downstream of GWAS and the genes within the significant windows of RHM were identified as candidate genes using the Ensembl BiomaRt R package [51, 52] based on the latest Bos taurus genome assembly of ARS-UCD1.3 and Bos taurus Annotation Release 113. Candidate genes were further validated by gene expression analyses using RNA-seq transcriptomic information between high and low DMI animal groups in the study population. High and Low DMI were selected based on the genomic breeding values (GEBV) of DMI for these 121 cows with both DMI information and RNA-seq transcriptomic profiles. The GEBV of DMI for these 121 cows were estimated and extracted from the UK national dairy genomic evaluation for feed efficiency [53]. Among the 121 cows, the 30 cows with the highest GEBV for DMI (i.e., High DMI group, mean GEBV(DMI) = 1.7392 kg/d) and the other 30 cows with the lowest GEBV for DMI (i.e., Low DMI group, mean GEBV(DMI) = -0.0004 kg/d) for each RNA-seq batch were compared for their gene expression between High and Low DMI groups for the DMI candidate gene identified by GWA and RHM analyses. The selected High and Low DMI groups (GEBV accuracies all higher than 0.6) were shown significantly differ in GEBV for DMI for both batches (p-values < 2.2e-16; Supplementary Figure 1).

We performed t-tests to compare log2(TPM+1) levels of gene expression for both batches of RNA-seq data for each potential causal gene for each lactation stage between the high 30 and low 30 GEBV(DMI) animal groups to compare the gene expression for candidate genes.

## Results

### GWAS signals for DMI

Summary statistics for SNPs that surpassed the genome-wide suggestive threshold are summarized in Table 2. The lambda estimates showed a slight inflation, and so SNP p-values were adjusted accordingly [44].

**Table 2.**
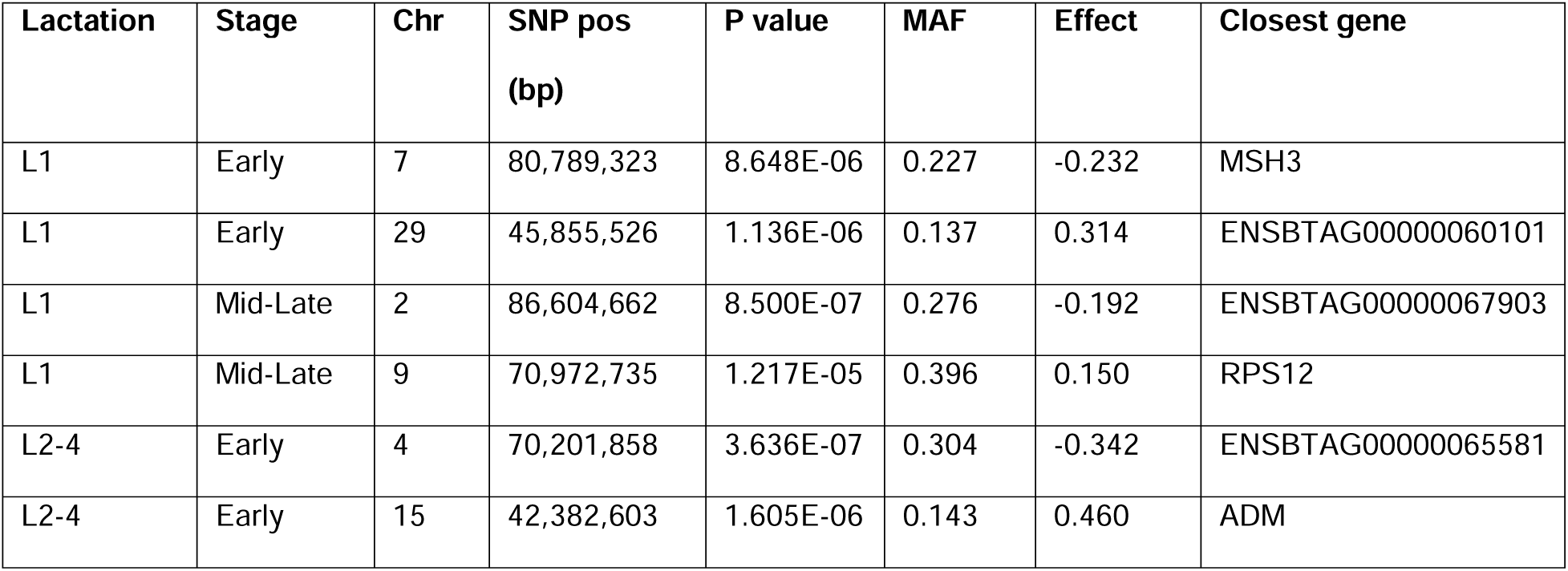
Summary statistics for SNPs that surpassed the genome-wide suggestive threshold for each trait from single-marker GWAS for Dry Matter Intake (DMI).

For early lactation 1, no SNPs passed the genome-wide significance threshold, two SNPs passed the genome-wide suggestive threshold; one SNP on chromosome 7 and one SNP on chromosome 29 (Figure 2A). For mid-late lactation 1, seven SNPs on chromosome 2 and one SNP on chromosome 9 passed the suggestive threshold (Figure 2B).

**Figure 2.**
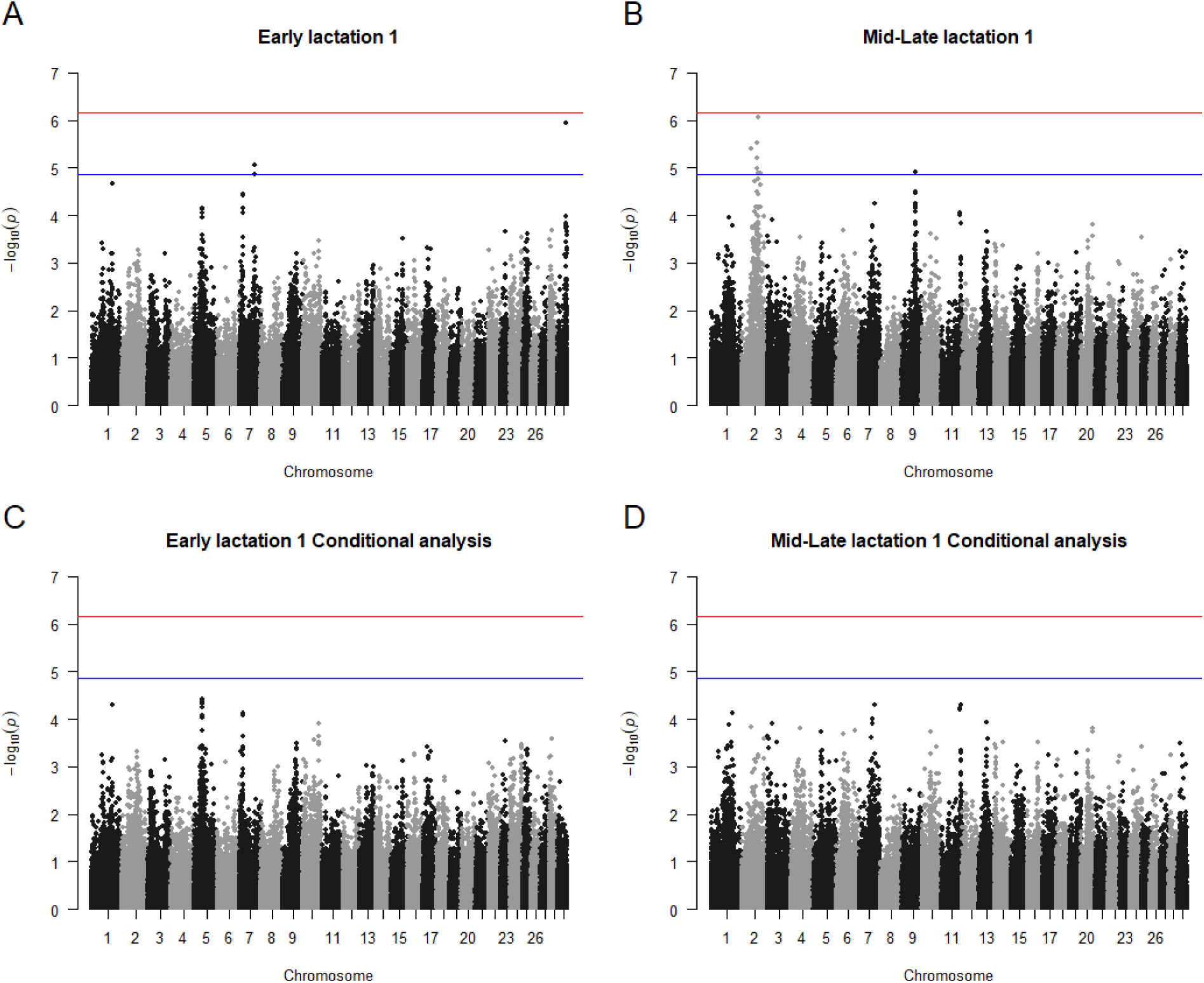
Manhattan plots for A) lactation 1, weeks 1-8 (early), B) lactation 1 weeks 9-44 (mid-late), C) lactation 1, weeks 1-8 after conditional analysis, and D) lactation 1 weeks 9-44 after conditional analysis. The blue line represents the genome-wide suggestive threshold, whilst the red line represents the genome-wide significance threshold.

For early lactations 2-4, one SNP on chromosome 4 passed the genome-wide significance threshold whilst another four SNPs on chromosome 4 and three SNPs on chromosome 15 passed the suggestive threshold (Figure 3A). No SNPs were suggestive for the mid-late stage (Figure 3B).

**Figure 3.**
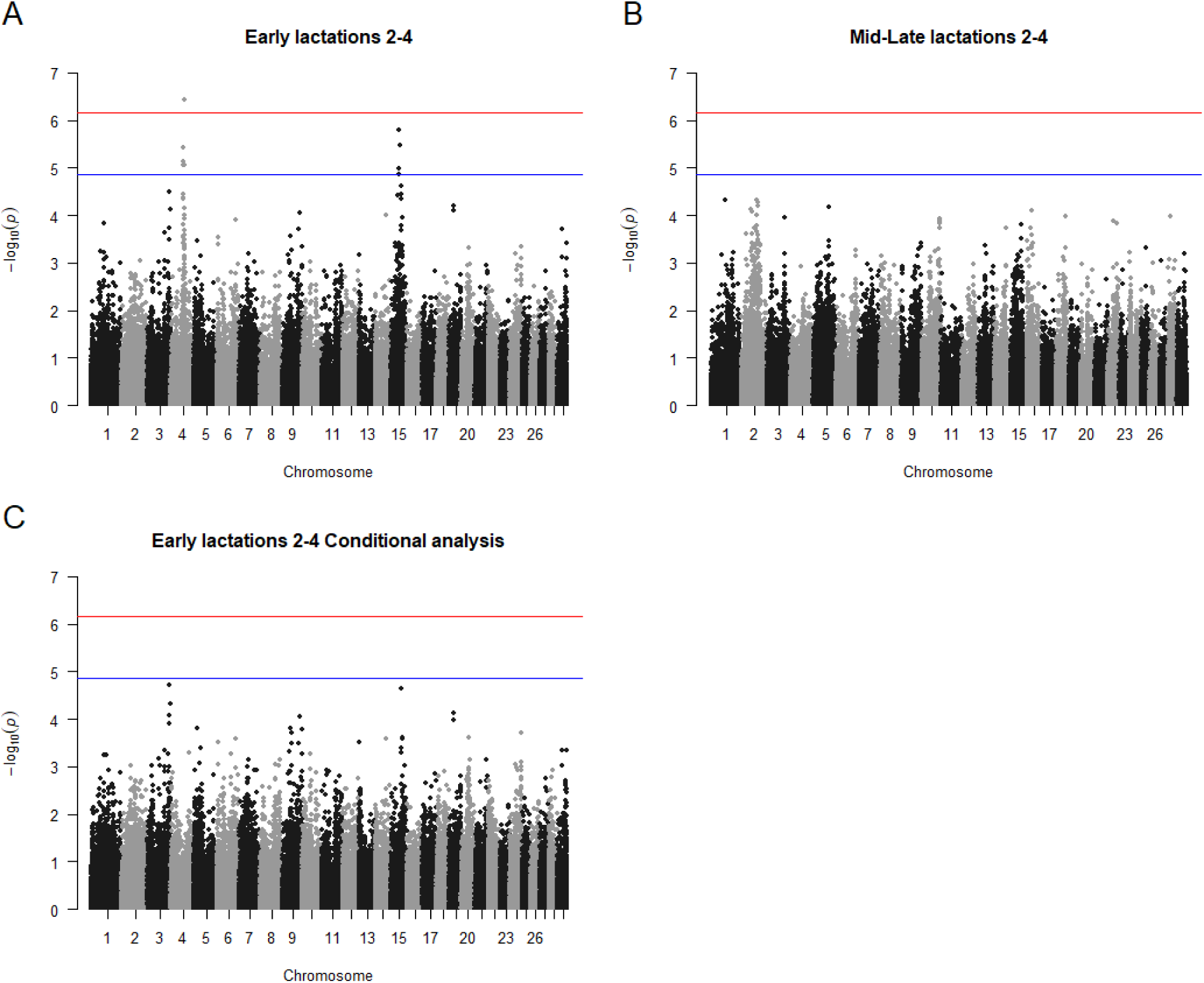
Manhattan plots for A) lactations 2-4, weeks 1-8 (early), B) lactations 2-4 weeks 9-44 (mid-late) and C) lactations 2-4 weeks 1-8 after conditional analysis. The blue line represents the genome-wide suggestive threshold, whilst the red line represents the genome-wide significance threshold. Conditional analysis was not performed for lactations 2-4 weeks 9-44 as no SNPs surpassed the genome-wide suggestive threshold for this lactation stage.

Conditional analyses of the chromosome-specific SNPs with the smallest p-values above the suggestive threshold showed that none remained suggestive, indicating the observed peaks do not represent multiple independent associations (Figures 2C, 2D, 3C and 3D).

### Significant genomic windows for DMI by RHM

When performing RHM for early lactation 1, ten 50-SNP windows surpassed the genome-wide significance threshold; two on chromosome 5, four on chromosome 7, one on chromosome 25 and three on chromosome 29. When reduced to fine-map 10-SNP windows within the significant 50-snp windows, six 10-snp windows were shown to be independently significant after conditional analysis: three on chromosome 7, two on chromosome 25 and one on chromosome 29 (Table 3).

**Table 3.** Significant independent 10-SNP genomic windows associated with Dry Matter Intake (DMI) from Regional Heritability Mapping (RHM). All 10-SNP windows surpass the significance thresholds.

| Lactation | Stage | Chr | Window start<br>bp | Window end<br>bp | P-value | Percentage of<br>genetic variance<br>explained | Significance<br>threshold |
| --- | --- | --- | --- | --- | --- | --- | --- |
| L1 | Early | 7 | 16,025,214 | 16,269,013 | 5.510E-04 | 0.030 (0.024) | 7.042E-04 |
| L1 | Early | 7 | 16,488,537 | 16,824,008 | 6.695E-06 | 0.025 (0.018) | 7.042E-04 |
| L1 | Early | 7 | 80,964,555 | 81,215,809 | 2.805E-05 | 0.053 (0.035) | 7.042E-04 |
| L1 | Early | 25 | 25,163,991 | 25,372,117 | 2.918E-07 | 0.031 (0.018) | 7.042E-04 |
| L1 | Early | 25 | 25,408,570 | 25,616,624 | 1.851E-05 | 0.007 (0.007) | 7.042E-04 |
| L1 | Early | 29 | 46,314,080 | 46,469,823 | 6.666E-04 | 0.028 (0.020) | 7.042E-04 |
| L1 | Late | 1 | 114,188,331 | 114,333,044 | 8.203E-08 | 0.015 (0.014) | 2.439E-04 |
| L1 | Late | 2 | 68,136,826 | 68,476,007 | 9.111E-05 | 0.096 (0.058) | 2.439E-04 |
| L1 | Late | 2 | 86,469,285 | 86,799,746 | 1.161E-13 | 0.065 (0.036) | 2.439E-04 |
| L1 | Late | 4 | 38,087,067 | 38,577,587 | 8.297E-08 | 0.015 (0.011) | 2.439E-04 |
| L1 | Late | 9 | 68,655,445 | 68,988,506 | 4.422E-09 | 0.019 (0.015) | 2.439E-04 |
| L2-4 | Early | 4 | 67,099,154 | 67,325,499 | 5.010E-04 | 0.005 (0.008) | 7.463E-04 |
| L2-4 | Early | 4 | 69,894,433 | 70,161,544 | 1.402E-06 | 0.041 (0.029) | 7.463E-04 |
| L2-4 | Early | 15 | 45,372,391 | 45,610,637 | 7.927E-06 | 0.059 (0.045) | 7.463E-04 |
| L2-4 | Late | 2 | 79,725,743 | 79,923,269 | 9.730E-08 | 0.018 (0.015) | 2.941E-03 |

For mid-late lactation 1, 31 50-SNP windows surpassed the genome-wide significance threshold; one on chromosome 1, 20 on chromosome 2, three on chromosome four and seven on chromosome 9. After conditional analysis, five 10-SNP windows independently surpassed the significance threshold: one on chromosome 1, two on chromosome 2, one on chromosome 4 and one on chromosome 9 (Table 3).

For early lactations 2-4, nine 50-SNP windows were significant; three on chromosome 4 and six on chromosome 15. Two 10-SNP windows on chromosome 4 and one on chromosome 15 were independently significant after conditional analysis (Table 3).

Three 50-SNP windows were significant for late lactations 2-4, all on chromosome 2. Conditional analysis showed that only one 10-SNP window on chromosome 2 was independently significant (Table 3).

### Gene expression validation of DMI candidate genes

In total, there were 45 genes identified as potential causal genes for feed intake in dairy cattle from the GWAS and RHM analyses; six from the GWAS results and 40 from the RHM (one gene was identified by both methods) (Table 4, Supplementary Figures 1 and 2). All these genes were analysed through gene expression analyses to compare their difference in gene expression between high and low DMI cow groups.

**Table 4.** Summary of the candidate genes for Dry Matter Intake (DMI) identified via GWAS and/or RHM, and whether they show significant difference in gene expression between individuals with high and low genomic breeding values of DMI.

| Lactation | Stage | Gene | Chr | GWAS | RHM | RNA-seq |
| --- | --- | --- | --- | --- | --- | --- |
| L1 | Early | FBN3 | 7 |  | ? |  |
| L1 | Early | ELAVL1 | 7 |  | ? |  |
| L1 | Early | TRAPPC5 | 7 |  | ? |  |
| L1 | Early | FCER2 | 7 |  | ? | ? |
| L1 | Early | EVI5L | 7 |  | ? | ? |
| L1 | Early | LRRC8E | 7 |  | ? |  |
| L1 | Early | MCEMP1 | 7 |  | ? | ? |
| L1 | Early | RETN | 7 |  | ? |  |
| L1 | Early | CCL25 | 7 |  | ? |  |
| L1 | Early | CD209 | 7 |  | ? |  |
| L1 | Early | MSH3 | 7 | ? |  |  |
| L1 | Early | STXBP2 | 7 |  | ? |  |
| L1 | Early | MAP2K7 | 7 |  | ? |  |
| L1 | Early | INSR | 7 |  | ? |  |
| L1 | Early | CLEC4G | 7 |  | ? |  |
| L1 | Early | SNAPC2 | 7 |  | ? |  |
| L1 | Early | TIMM44 | 7 |  | ? |  |
| L1 | Early | RASGRF2 | 7 |  | ? | ? |
| L1 | Early | ARHGEF18 | 7 |  | ? |  |
| L1 | Early | PRR36 | 7 |  | ? |  |
| L1 | Early | ENSBTAG00000046336 (novel gene) | 7 |  | ? |  |
| L1 | Early | ENSBTAG00000047612 (novel gene) | 7 |  | ? |  |
| L1 | Early | ENSBTAG00000069568 (novel gene) | 7 |  | ? |  |
| L1 | Early | GSG1L | 25 |  | ? |  |
| L1 | Early | KATNIP | 25 |  | ? |  |
| L1 | Early | TPCN2 | 29 |  | ? |  |
| L1 | Early | ENSBTAG00000046090 (novel gene) | 29 |  | ? |  |
| L1 | Early | ENSBTAG00000060101 (novel gene) | 29 | ? |  |  |
| L1 | Early | ENSBTAG00000062766 (novel gene) | 29 |  | ? |  |
| L1 | Early | MRGPRF | 29 |  | ? |  |
| L1 | Mid-late | PLCL1 | 2 |  | ? | ? |
| L1 | Mid-late | ENSBTAG00000062557 (novel gene) | 2 |  | ? |  |
| L1 | Mid-late | ENSBTAG00000067903 (novel gene) | 2 | ? | ? |  |
| L1 | Mid-late | CACNA2D1 | 4 |  | ? |  |
| L1 | Mid-late | RPS12 | 9 | ? |  |  |
| L1 | Mid-late | EPB41L2 | 9 |  | ? |  |
| L1 | Mid-late | ENSBTAG00000059877 (novel gene) | 9 |  | ? |  |
| L2-4 | Early | CPVL | 4 |  | ? |  |
| L2-4 | Early | CREB5 | 4 |  | ? |  |
| L2-4 | Early | TRIL | 4 |  | ? |  |
| L2-4 | Early | ENSBTAG00000058735 (novel gene) | 4 |  | ? |  |
| L2-4 | Early | ENSBTAG00000065581 (novel gene) | 4 | ? |  |  |
| L2-4 | Early | ADM | 15 | ? |  | ? |
| L2-4 | Early | SYT9 | 15 |  | ? |  |
| L2-4 | Mid-late | MYO1B | 2 |  | ? |  |

In early lactation 1, four candidate genes (*FCER2*, *EVI5L*, *MCEMP1* and *RASGRF2*) identified by GWAS and RHM also showed significant differences in gene expression between the 30 individuals with the highest DMI GEBV and the 30 individuals with the lowest DMI GEBV in at least one of the RNA-seq batches (Figure 4, Table 4). In mid-late lactation 1, one gene (*PLCL1)* identified by GWAS and RHM also showed significant differences in gene expression between individuals with the highest and lowest DMI GEBV in at least one of the RNA-seq batches (Figure 4, Table 4).

**Figure 4.**
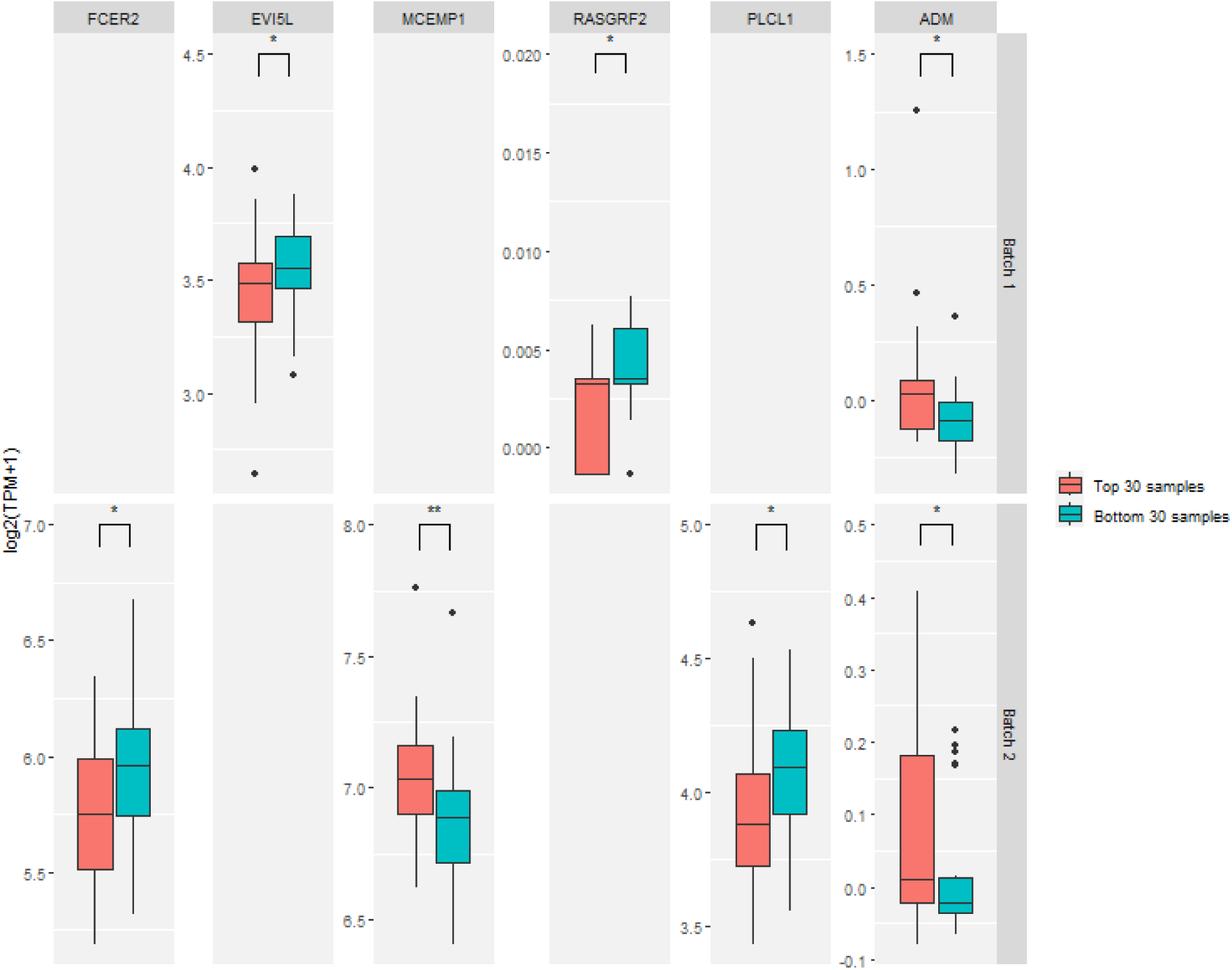
Comparison of gene expression levels for genes identified via GWAS and RHM that showed significant difference in expression between individuals with high and low DMI GEBV. Red indicates individuals with high GEBV, blue indicates individuals with low GEBV. * = p value < 0.05, ** = p value < 0.005

In early lactations 2-4, one gene (*ADM*) identified by GWAS and RHM also showed significant differences in expression between individuals with the highest and lowest DMI GEBV in both RNA-seq batches (Figure 4, Table 4). One gene was found to be associated with mid-late lactations 2-4 (*MYO1B*), however it did not show significant differences in expression in either of the two RNA-seq batches (Figure 4, Table 4).

## Discussion

In this study, our GWAS and RHM results indicated clear differences in genetic architecture underlying DMI both between primiparous and multiparous cows, and between early and mid-late lactation stages within lactation. Similar findings have been seen by [28] showing that different genetic variants contribute to DMI in lactating Holstein cattle depending on how many days post-partum an individual is. In addition, the difference in genetic basis for DMI between lactation stages were also found by [21, 28] where they showed low genetic correlations between DMI in early lactation and in mid-late lactation stages. Considering the accumulated evidence of the change of genetic basis for dairy feed intake between lactations (mainly primiparous vs. multiparous cows) and between lactation stages (mainly early vs. mid-late lactation stage), feed intake from different lactations/stages may need to be treated as separate traits in genetic and genomic analyses.

Using both GWAS and RHM, we have identified 45 potential causal genes associated with DMI during different stages of lactation in primiparous and multiparous cattle, six of which also showed significant differences in gene expression between individuals with high and low DMI GEBV. Compared with GWAS, Regional Heritability Mapping in this study enabled an increased power to detect regions containing multiple alleles that individually contribute too little variance to be detectable by GWAS as well as regions with single common GWAS-detectable SNPs [33–36]. The use of transcriptomic information in this study facilitated the validation of candidate genes identified by solely the genomic analyses, offering extra information to validate functional variants using multi-omics data.

Six candidate genes were identified by GWAS/RHM and were further validated by gene expression analyses, which are of high interest for DMI. Three of these genes have known links to sugar and adipose metabolism, milk production, and body weight:

*ADM*, *EVI5L* and *PLCL1*. The *ADM* gene, associated with DMI in early lactation of multiparous cows, has been shown to be involved in insulin signalling [55] and is found to have increased expression in the adipose tissues of obese individuals [54]. The *EVI5L* gene, associated with DMI in early lactation 1, has previously been found to be involved in milk production in goats [55]. The *PLCL1* gene, associated with late lactation 1, has previously been found to be associated with weight in lambs between months 3-7 of life [56]. In addition, two candidate genes, associated with DMI in early lactation 1, have a link to immune function: *MCEMP1* and *FCER2*. *MCEMP1* is expressed on the membrane of mast cells and monocytes [57] and is involved in inflammatory immune response [58]. And *FCER2* is an IgE receptor [59] and is involved in immune response to parasites [60] and allergies [61]. Finally, RASGRF2 – the sixth gene that showed significant differences in expression between high and low DMI individuals – has been shown to be associated with levels of cocaine self-administration in mice [62] and binge-drinking in humans [63], suggesting a role in the dopamine reward pathway. The dopamine reward pathway is known to modulate the “rewarding” sensation of food and is thought to play a role in obesity in humans [64] – this finding suggests that a similar effect may occur in cattle, influencing feed intake. This gene is associated with early lactation 1.

It is possible that some of the candidate genes we identified that did not show a significant difference in blood gene expression levels between high and low DMI individuals may still play a role in DMI variation, but through a different tissue other than blood. Our study focused on gene expression in blood as blood sampling is non-invasive and DNA biomarkers in blood is the easiest to collect for future large-scale breeding purposes. However, blood may not fully capture the transcriptional dynamics relevant to DMI as the physiological processes governing feed intake occur in multiple tissues along the digestive tract and are likely influenced by neural regulation. Therefore, future studies using gene expression data from multiple tissues may provide further evidence of the DMI candidate genes identified in this study.

## Conclusions

This study highlighted the complex genetic architecture underlying dairy feed intake across lactations and lactation stages, where clear differences in the genetic architecture were observed not only between lactations but also between stages within lactation, reflecting dynamic changes in the physiological and metabolic demands of dairy cattle. This variation highlights the need for high-resolution data analysis to capture these subtle but critical differences that can inform genetic and management strategies for optimizing DMI and overall productivity. Through genomic and transcriptomic analyses, candidate genes for DMI were highly linked to sugar and adipose metabolism, milk production, body weight, dopamine-reward pathways and immune functions.

## List of abbreviations

DMI: Dry matter intake
SNP: Single nucleotide polymorphism
GWAS: Genome-wide association study
RHM: Regional Heritability Mapping
QTL: Quantitative trait loci
DM%: Dry matter content
LRT: Log likelihood ratio test
GRM: Genomic relationship matrix
GEBV: Genomic breeding values

## Declarations

### Ethics approval and consent to participate

Not applicable

### Consent for publication

All authors have read and approved the final manuscript.

### Availability of data and materials

The GWAS summary statistics and the RNA-seq data are available by request. Scripts used for this analysis are available at https://github.com/CaelinnJames/GenomicAndTranscriptomicAnalysesofFI

## Authors’ contributions

CJ performed the analyses and wrote the manuscript. BL secured the funding and designed the experiment, collected the samples, contributed to writing of the manuscript. LF contributed to blood sampling and RNA-seq data analyses. ZW and JH contributed to the RNA extraction of the blood samples for sequencing. MC contributed to the phenotype collection and the genotype information of the study. All authors read and approved the final manuscript.

## Acknowledgements

The authors acknowledge Research Computing at the James Hutton Institute for providing computational resources and technical support for the “UK’s Crop Diversity Bioinformatics HPC” (BBSRC grants BB/S019669/1 and BB/X019683/1), use of which has contributed to the results reported within this paper. All computational analyses described in this paper were performed on Crop Diversity HPC [65]. Ian Archibald (SRUC, Edinburgh, UK) is acknowledged for curating the Langhill data, Jenni Flockhart (SRUC) for details of the Langhill diets, and all staff at Crichton farm for assisting sample collection.

---

**Supplementary Figure 1.**
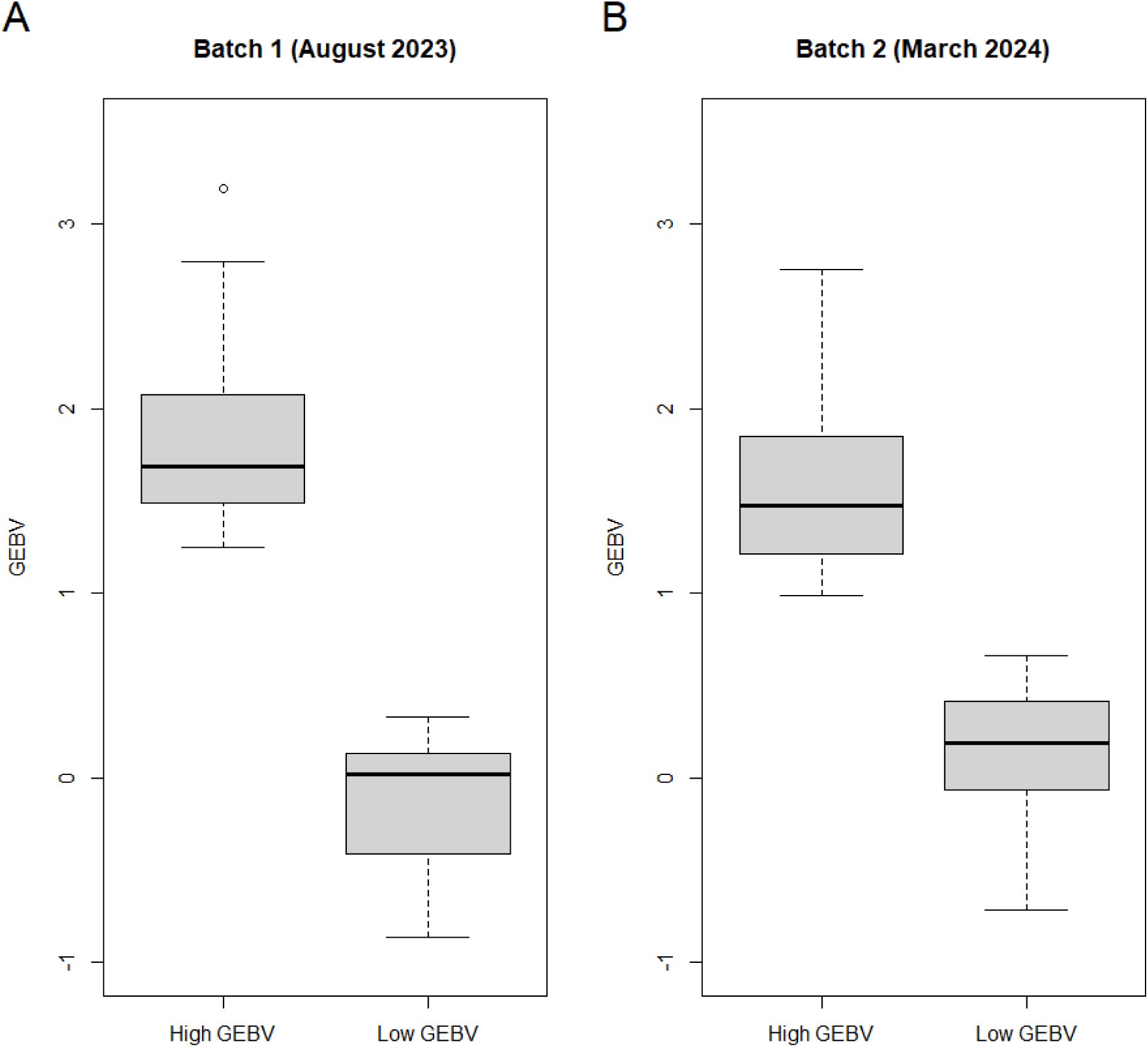
Comparison of DMI GEBV for the 30 individuals with the highest GEBV and 30 individuals with the lowest GEBV for all individuals sampled in August 2023 (A) and all individuals sampled in March 2024 (B).

**Supplementary Figure 2.**
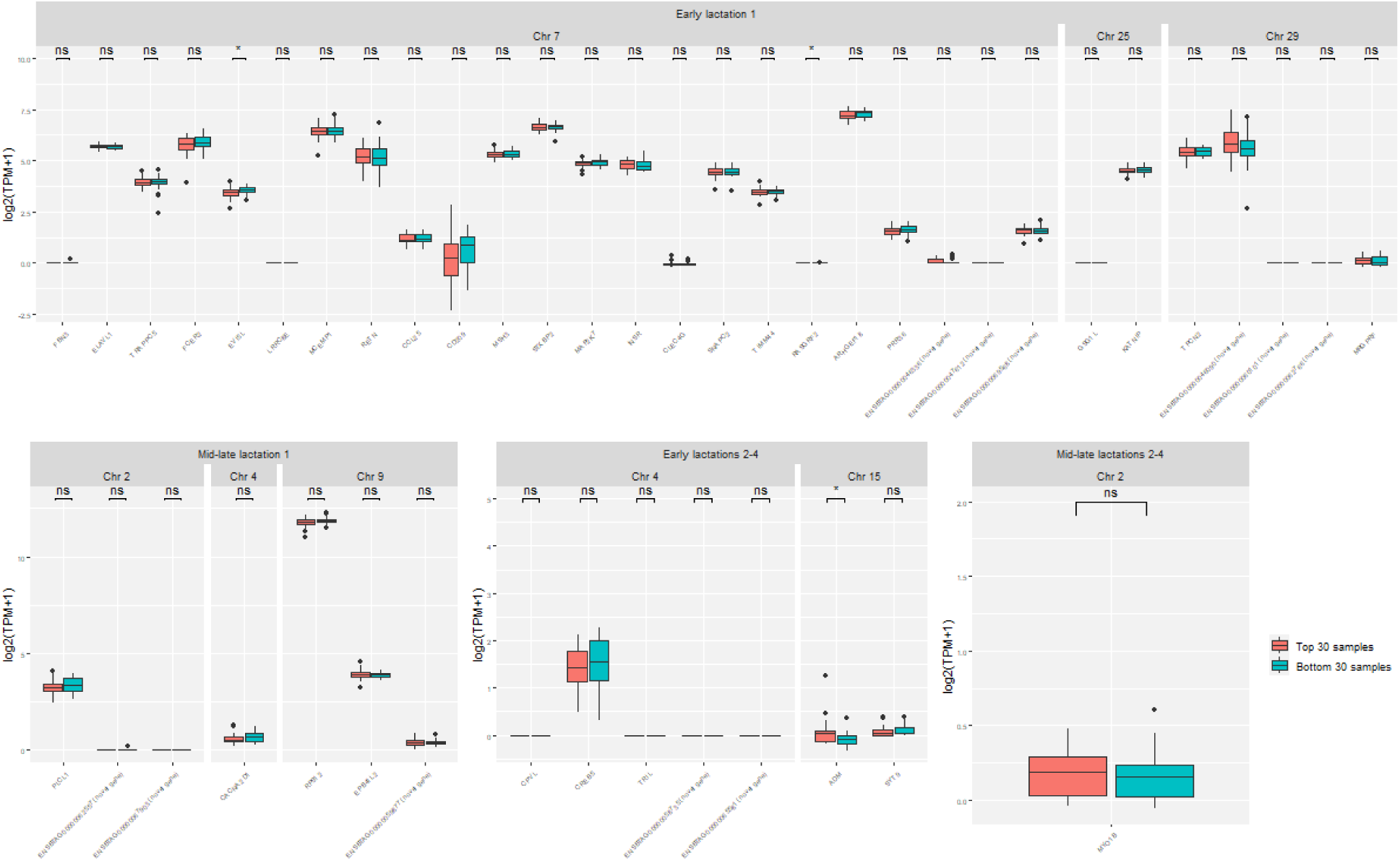
Comparison of gene expression levels for genes identified via GWAS and RHM for individuals with high and low DMI GEBV for individuals sampled in August 2023 (batch 1). Red indicates individuals with high GEBV, blue indicates individuals with low GEBV. Ns= p value > 0.05 * = p value < 0.05, ** = p value < 0.005

**Supplementary Figure 3.**
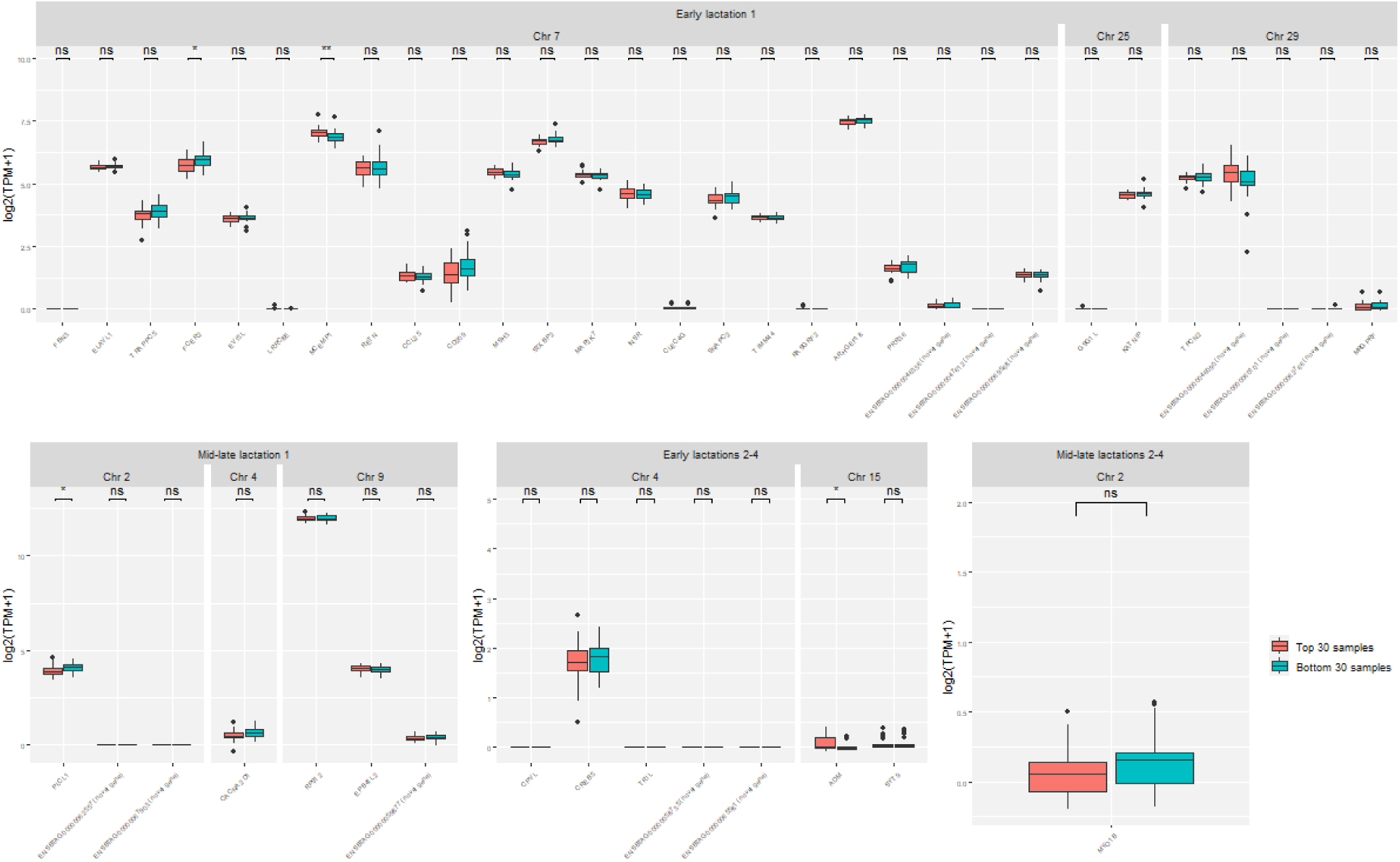
Comparison of gene expression levels for genes identified via GWAS and RHM for individuals with high and low DMI GEBV for individuals sampled in March 2024 (batch 2). Red indicates individuals with high GEBV, blue indicates individuals with low GEBV. Ns= p value > 0.05 * = p value < 0.05, ** = p value < 0.005

